# Mapping Patient Trajectories using Longitudinal Extraction and Deep Learning in the MIMIC-III Critical Care Database

**DOI:** 10.1101/177428

**Authors:** Brett K. Beaulieu-Jones, Patryk Orzechowski, Jason H. Moore

## Abstract

Electronic Health Records (EHRs) contain a wealth of patient data useful to biomedical researchers. At present, both the extraction of data and methods for analyses are frequently designed to work with a single snapshot of a patient’s record. Health care providers often perform and record actions in small batches over time. By extracting these care events, a sequence can be formed providing a trajectory for a patient’s interactions with the health care system. These care events also offer a basic heuristic for the level of attention a patient receives from health care providers. We show that is possible to learn meaningful embeddings from these care events using two deep learning techniques, unsupervised autoencoders and long short-term memory networks. We compare these methods to traditional machine learning methods which require a point in time snapshot to be extracted from an EHR.

## 1. Introduction

After the U.S. government mandated meaningful use of electronic health records (EHRs) by 2014, they have been widely adopted with 96% of health care providers implementing an EHR [1]. Patient interactions with the health care system are recorded in the EHR. Many research analyses treat the EHR as a static document by taking a snapshot of a patient’s EHR and using this for downstream analyses. This fails to account for the way a patient changes over time, their trajectory.

Jensen et al. [2] proposed the idea of temporal disease trajectories to model expected progression for a patient over time. This study uses billing codes as disease labels, which may introduce biases inherent to the billing process. Patients may be assigned a billing code before being diagnosed for a disease in order to receive a diagnostic test. Billing codes place also binary rules on the presence of disease. Perhaps most importantly for this work billing codes are frequently assigned after a visit and are thus not helpful for tracking patient trajectories over the course of an inpatient admission or rapid series of visits.

Interactions between patients and the health care system tend to occur in bursts, related to a specific visit or a series of visits. We label these periods of activity as care events and group these actions together. These care events represent changes over time and can capture longitudinal changes of a patient’s state.

Denny et al. [3] first showed the ability to use autoencoders to model clinical measures in an unsupervised manner. More recently, several groups have used autoencoders to learn high level features useful for classification [4,5] and imputation [6]. Tan et al. also showed the ability to extract meaningful features from gene expression data using autoencoders [7]. We use autoencoders to represent patient care events in a low dimensional vector space that is useful for visualization. Positions in this vector space represent the patient’s condition at a point in time. By connecting these positions, or care events, in order, it is possible to see how a patient’s condition changes over time and how they move through the health system. It is also possible to cluster patients in this low dimensional space and examine when patient outcomes diverge, one group having high survival and the other having high mortality.

This care event representation also provides a natural sequence of events. Recurrent neural networks have shown an impressive ability to model sequences to solve problems in many domains including object recognition in computer vision [8], image [9] and text generation [10]. Long short-term memory networks (LSTMs) [11] are a type of recurrent neural network that have recently been applied to clinical data to learn low dimension representations of medical concepts [12] and to make classifications using time series of specific clinical measures [13,14]. Trajectories have been used to model multistage dynamic decision processes (DMP) in discrete optimization problems [15]. In Algebraic Logical Meta-Model (ALMM) the state of the system in a certain time depends on the previous state, undertaken decision and transition function. This concept allows to easily describe the state of the patient at a particular time, with specific actions taken (e.g. application of medication) to manage the response to previous events within the progression of a disease.

In this work, we first demonstrate that deep learning approaches can (1) learn patient embeddings useful for both interpretable expert analysis via visualization and (2) do this we use the Medical Information Mart for Intensive Care III (MIMIC) database and apply both unsupervised deep autoencoders and LSTMs.

## 2. Methods

### 2.1. Source Code and Analysis Availability

Source code to reproduce the analyses in this work are provided in our repository (https://github.com/EpistasisLab/MIMIC_trajectories) under a permissive open source license. In addition, Continuous Analysis [16] was used to generate docker images matching the environment of the original analysis.

### 2.2. Care Event Extraction

#### 2.2.1. Medical Information Mart for Intensive Care III (MIMIC) Critical Care Database

MIMIC [17] is a publicly available database composed of 46,297 critical care de-identified electronic health records for patients at Beth Israel Deaconess Medical Center. It includes all charted data (demographics, vital signs, medications, procedures, diagnoses, patient outputs, laboratory tests, physician notes, and treatment details) for patients from 2001 to 2012.

#### 2.2.2. Extracting Care Events from MIMIC

We divided the MIMIC database into 4 groups:

1. Static data that does not change over the course of an admission (i.e. demographic data).
2. Actions performed by health care providers that have a specific time associated with them (i.e. laboratory events).
3. Actions performed by health care providers that only have a date associated with them (i.e. oral medications).
4. Streaming data measured on a per-minute basis (i.e. heart rate).

To define care events, we included all actions initiated, or charted, by health care providers that have a specific time associated with them (Table 1). These actions were placed in sequential order and grouped together until there was a gap greater than the margin time (Figure 1). Because this is critical care data, the timeline between events is much smaller than typical EHR data. We found a 59 minute margin time yielded care events that had a good balance of inclusiveness while not including extended time periods. This yielded 1,566,026 total care events and an average of 26.80 care events per admission. In outpatient datasets, we expect a margin time of several days may better capture the concept of a care event.

**Table 1.**
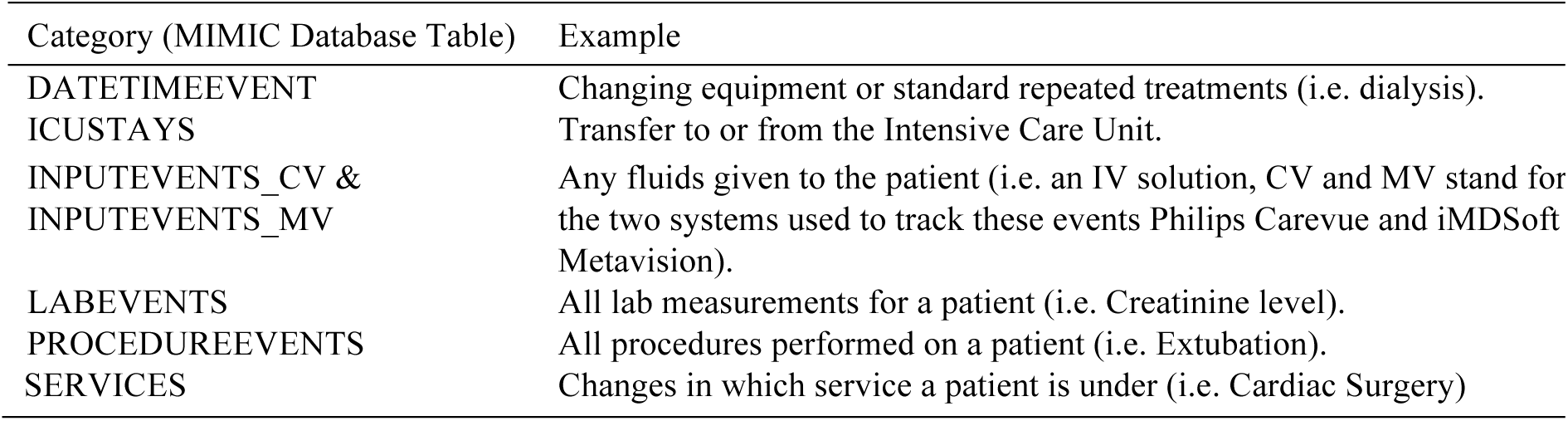
*Categories and examples of Care Event Actions.*

| Category (MIMIC Database Table) | Example |
| --- | --- |
| DATETIMEEVENT | Changing equipment or standard repeated treatments (i.e. dialysis). |
| ICUSTAYS | Transfer to or from the Intensive Care Unit. |
| INPUTEVENTS_CV &<br>INPUTEVENTS_MV | Any fluids given to the patient (i.e. an IV solution, CV and MV stand for the two systems used to track these events Philips Carevue and iMDSof Metavision). |
| LABEVENTS | All lab measurements for a patient (i.e. Creatinine level). |
| PROCEDUREEVENTS | All procedures performed on a patient (i.e. Extubation). |
| SERVICES | Changes in which service a patient is under (i.e. Cardiac Surgery) |

**Fig. 1.**
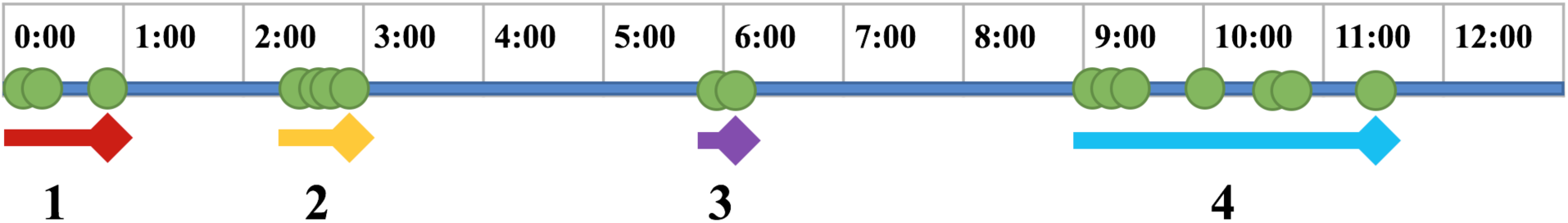
Example of care event extraction. Green circles indicate actions taken by health care providers. Lines and numbers below indicate care events.

#### 2.2.3. Stratification of Patient Attention based on type of Insurance Provider

Care events can provide a useful heuristic to the level of interaction between the health care provider and a patient. To evaluate attention, we compared the time spent in the hospital per admission with the number of care events per admission and the average number of care events per day. We then performed Welch’s t-test between patients with private insurance and each of the other types of insurance (Medicare, Medicaid, Government, Self-Payment) to see if there were significant differences between patients with differing insurance types.

### 2.3. Unsupervised learning to learn embeddings of extracted Care Events

#### 2.3.1. Applying Autoencoders to Extracted Care Events to cluster in a low dimensional space

We used the Keras library [18] to construct autoencoders with 7 hidden layers in (1196, 512, 256, 128, 64, 128, 256, and 512 nodes per layer). We used dropout to mask 20% of the connections between the input layer and the first hidden layer. The model was trained using binary cross entropy loss with Adam [19]. The middle, hidden layer (64 nodes) was used as an output for visualization using t-Stochastic Neighbor Embedding [20]. The resulting visualizations were labeled for enrichment of 1-year patient survival. Survival data was based on the date of death variable in the MIMIC dataset, a merger between the hospital and social security data.

### 2.4. Predicting Survival Using Care Events

We evaluated how effectively different machine learning methods could predict patient survival over a 1-year period (as measured from the original admission date). The 1-year survival period began on the date of admission.

For this analysis, we performed 5-fold cross validation providing a training set of 46,751 admissions and a test set of 11,687 admissions chosen via stratified cross validation [21]. Survival was predicted using several classifiers: (1) a standard feed forward or multi-layer perceptron deep neural network [18], (2) a random forest, (3) logistic regression and (4) support vector machine (linear kernel) [21] after various numbers (*N*) of care events: 1, 3, 5, 10, 20, 30 and 50. Area under the curve of the receiver operating characteristic was used for evaluation and comparison.

#### 2.4.1. Traditional machine learning methods to predict survival from an EHR Snapshot

To build a snapshot vector useable for traditional machine learning methods. We took the mean of each value from a set of care events, up to the *N*^th^ care event. If the patient had less than *N* care events, we took the mean for all of their care events. Feature selection was then performed on this aggregate vector using ReliefF [22,23] to choose the top 100 features. These 100 features were then provided as input to each of the machine learning classifiers.

#### 2.4.2. Long Short Term Memory Networks (LSTMs) to predict survival with Care Events Sequences

To build the sequence vector from a set of care events we first truncated sequences longer than *N*. Sequences shorter than N were post-padded with zeros. The model was comprised of 3 types of layers, an initial embedding layer, three LSTM layers (with 100, 50 and 50 nodes respectively) and a fully connected (Dense) output layer. We trained the model using rmsprop [24] with a binary cross entropy loss function.

## 3. Results

The MIMIC dataset includes 58,438 admissions from 46,297 unique patients. This was extracted to form 1,566,026 care events (Table 2). Medicare patients were double the age of other patients on average. Patients using private or government insurance and Medicaid had relatively equal mortalities during the initial admission and the next 6 months. Patients using Medicare had significantly higher mortality in the 6 months after admission as their time under critical care and self-payment patients had high mortality during the admission but lower admission after leaving critical care.

**Table 2.** Summary statistics for MIMIC Critical Care database.

|  | Total | Male | Female | Private | Medicare | Medicaid | Government | Self |
| --- | --- | --- | --- | --- | --- | --- | --- | --- |
| Patients | 46,297 | 26,121 | 20,399 | 19,663 | 21,002 | 4,570 | 1,614 | 600 |
| Admissions | 58,438 | 32,950 | 26,026 | 22,250 | 28,103 | 5,713 | 1,767 | 605 |
| Admissions per Patient | 1.26 | 1.26 | 1.28 | 1.13 | 1.34 | 1.25 | 1.09 | 1.01 |
| Average Age at Admission | 56.01 | 54.95 | 57.34 | 37.82 | 75.95 | 37.91 | 35.15 | 39.11 |
| Care Events | 1,566,026 | 867,941 | 698,085 | 637,968 | 693,254 | 179,182 | 46,722 | 8,900 |
| Care Events per Admission | 26.80 | 26.34 | 26.82 | 28.67 | 24.67 | 31.36 | 26.44 | 14.71 |
| Visit Survival | 90.84% | 90.38% | 89.56% | 95.24% | 86.40% | 94.60% | 95.87% | 85.2% |
| 6-Month Survival | 79.81% | 79.63% | 78.39% | 89.84% | 69.47% | 87.61% | 91.62% | 83.31% |
| 12-Month Survival | 76.28% | 76.19% | 74.82% | 87.90% | 64.33% | 84.75% | 90.32% | 82.81% |

### 3.1. Treatment and Outcome Comparison

We examined the length of stay per admission by insurance type (Figure 2A) and found that patients using Medicare had the longest stays but that all groups differed significantly via an ANOVA test (p-value 5.02E-28). In addition, we compared each type of insurance against the private group using Welch’s t-test. It is not surprising that patients using self-payment had the shortest stays and the least number of care events per stay (Figure 2B). Interestingly, patients with private insurance had significantly lower care events per day than the most similar (by age) other groups, government-based insurance and Medicaid (Figure 2B).

**Fig. 2.**
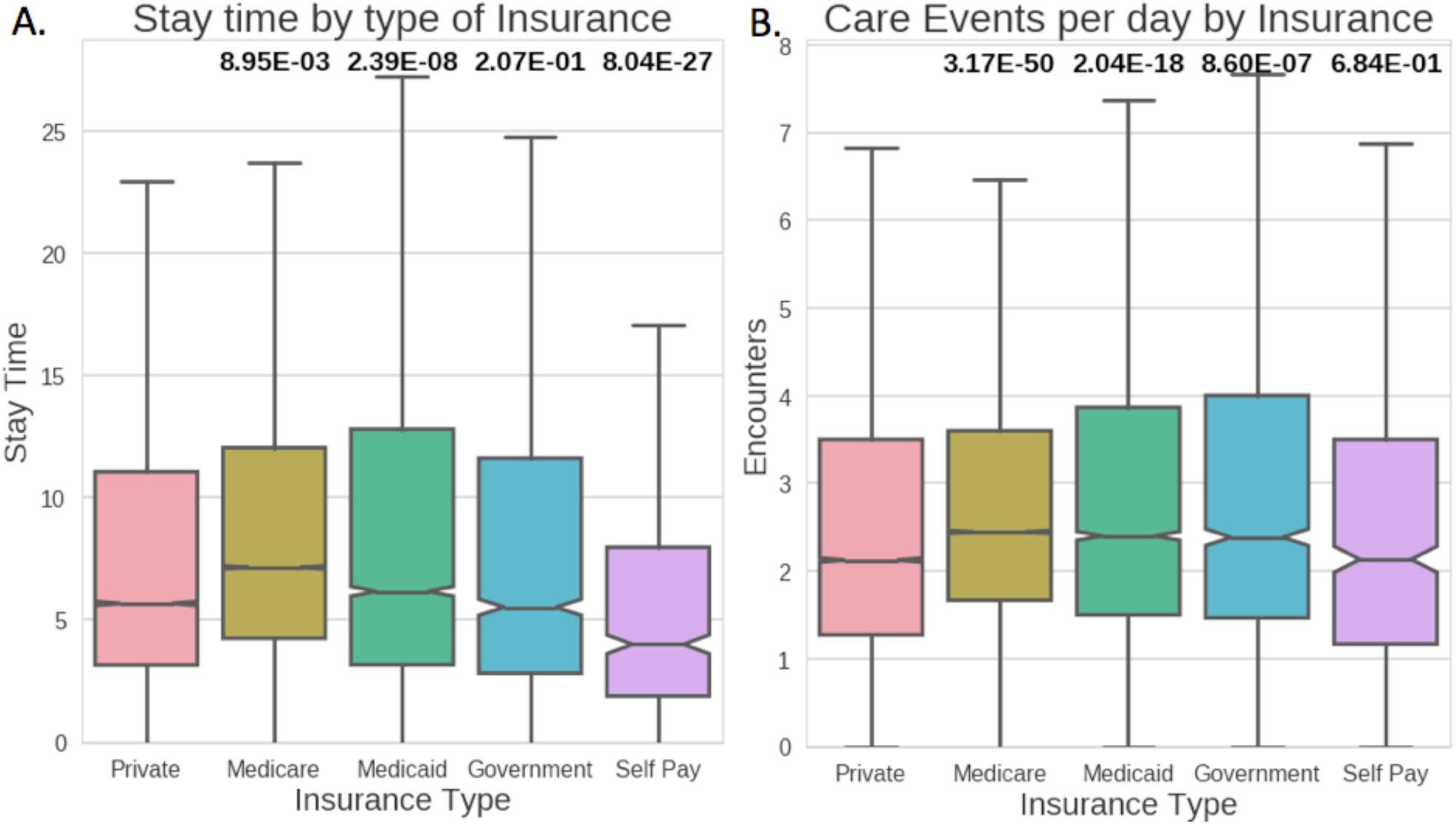
Association testing between different insurance types. A.) Length of admission. C) Number of care events per day of each admission. Labels at the top indicate p-values via Welch’s t-test to private group.

### 3.2. Unsupervised modeling of patient care events

To test whether unsupervised autoencoders could learn meaningful embeddings from individual care vents, we plotted the innermost hidden layer using t-Stochastic Neighbor Embedding (t-SNE) and overlaid 1-year survival labels (Figure 3). Figure 3 shows an unsupervised clustering, where the X and Y axes do not have an explicit meaning or interpretation. This clustering process yielded several clusters with high enrichment for either mortality or survival indicating the ability to learn meaningful embeddings. t-SNE does not maintain global similarity structure and as such this process is useful for visualizing single care events but not for understanding patient trajectories. To examine patient trajectories, it is necessary to look at the value of the innermost hidden layer before t-SNE was applied or to use a method designed to model sequential data. Recurrent neural networks, and specifically LSTMs are well suited at this task.

**Fig. 3.**
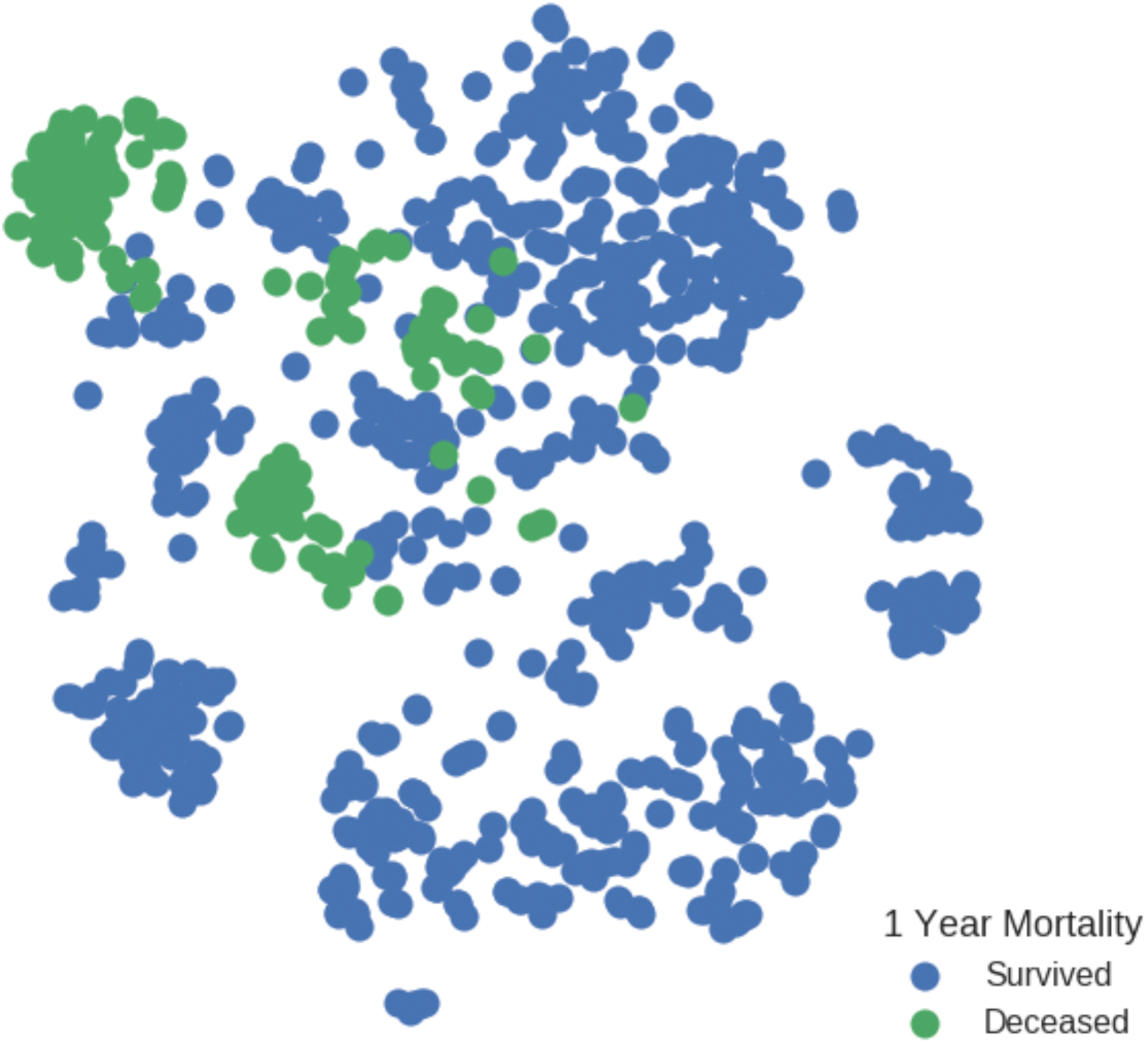
Unsupervised Care Event Embedding by applying t-SNE to the innermost layer of autoencoder (1000 care events shown to prevent overplotting).

### 3.3. Supervised prediction of patient survival

Next, we preformed the supervised classification task of predicting whether a patient survived one year from the date of their admission. We measured classification accuracy with differing numbers of care events to evaluate whether the care event-based approach had advantages over traditional single point in time measurements (Figure 4). The Pearson correlation of the number of care events to 1-year mortality rate was 0.062. Of the methods predicting based on a snapshot, the random forest was by far the most effective. Despite this, it did not increase in performance as more information about an admission was added. This indicates that much of its predictive power comes from the initial presentation. Both, linear methods and a traditional feed-forward neural network barely outperformed random chance. This may have been due to the high dimensionality of the dataset. The care event-based LSTM increases in performance as more care events are provided. This is particularly evident when more than the median number of care events (26.8) are provided as input to the LSTM. Including more than 50 care events yielded weaker results for the LSTM. This is likely because most patients have fewer than 50 care events so most of the signal is captured in the first 50 care events. Going beyond 50 leads to a high level of padding to signal.

**Fig. 4.**
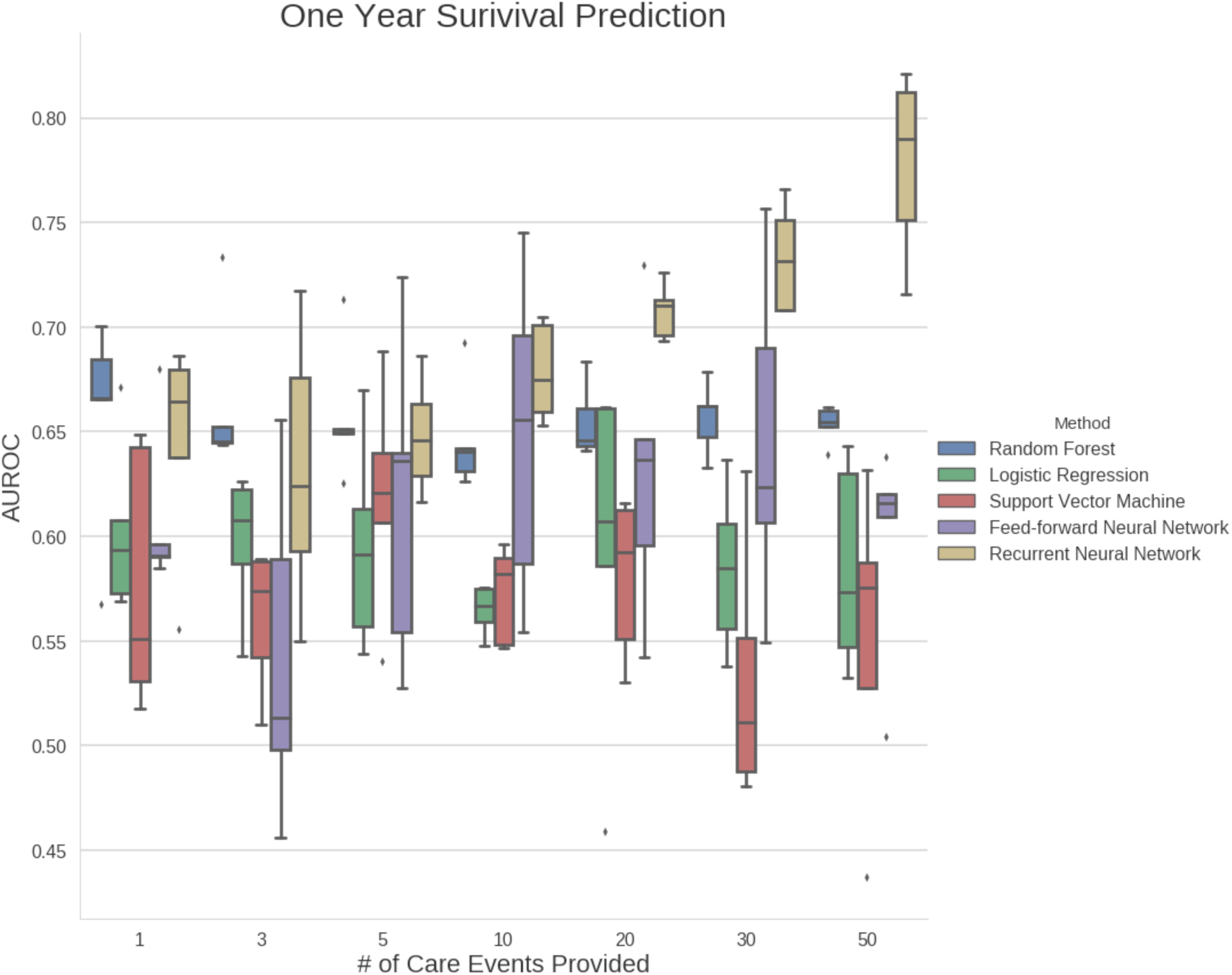
Comparison of machine learning methods and the number of care events provided for 1-year survival prediction (AUROC).

## 4. Discussion and Conclusions

By limiting the usage of summary statistics to small time periods, we offer a granular method for modeling longitudinal clinical data. The care event extraction method provides a simple data driven approach to extracting temporal data for use in time series analyses. It allows summary statistics to be computed over short time windows as opposed to an entire patient history or arbitrary timestamps. Care events also offer a heuristic to allow comparison of the level of attention different patients receive from health care providers. We demonstrated the ability to learn embeddings enriched for different endpoints using unsupervised deep learning and were able to more accurately predict patient survival using supervised long short-term memory networks.

Though our approach showed strong performance for several tasks in this dataset, this method currently has limitations in terms of generalization. Long-short term memory networks, like many deep learning approaches, require many patients to outperform other methods. This can present a challenge when studying a single phenotype instead of a wide variety of critical care patients. The greatest benefits are likely to be seen when patients have many care events, making this approach particularly well suited for chronic diseases like type 2 diabetes and Crohn’s disease or for diseases that are hard to subtype such as multiple sclerosis. An additional challenge is if a patient with a disease like type 2 diabetes suffers an unrelated acute injury (i.e. broken rib in a vehicle accident) this acute injury may introduce too much noise to capture the type 2 diabetes trajectory.

In future work, we hope to introduce filtering techniques to exclude or deemphasize unrelated diagnoses. We also plan to increase the dimensionality of the encoders and applying additional techniques of visual clustering [25]. This includes using Shared-Nearest Neighbors (SNN) clustering to find groups of patients with similar stage of the disease in noisy data and Mukres algorithm to map groups of patients resembling a state of the disease to clusters found in the data.

Another challenge we would like to take is including streaming data in the simulation. Some measurements, e.g. heart rate or blood pressure, are performed every minute for each patient. The information about sudden changes of patient’s condition is especially relevant for intensive-care patients. While our method aggregates patient data over shorter time periods than are commonly used, we plan to adapt our model by adding more detailed relevant information extracted from streaming sources.

## 5. Acknowledgments

We thank Casey S. Greene (University of Pennsylvania), Daniel S. Herman (University of Pennsylvania) and Andrew L. Beam (Harvard Medical School) for their helpful discussions. Funding: This work was supported by the Commonwealth Universal Research Enhancement (CURE) Program grant from the Pennsylvania Department of Health. B.K.B.-J., P.O. and J.H.M. were also supported by US National Institutes of Health grants AI116794 and LM010098 to J.H.M. Author Contributions: B.K.B.-J. and J.H.M. conceived of the study. B.K.B.-J. and P.O. performed initial data processing. B.K.B.-J. performed analyses and wrote the manuscript. All authors revised and approved the final manuscript. Competing Interests: The authors have no competing interests to disclose. Source code availability: All source code is available via github (https://github.com/epistasislab/MIMIC_trajectory).

